# The kinesin KIF4 mediates HBV/HDV entry through regulation of surface NTCP localization and can be targeted by RXR agonists *in vitro*

**DOI:** 10.1101/2021.09.29.462331

**Authors:** Sameh A. Gad, Masaya Sugiyama, Masataka Tsuge, Kosho Wakae, Kento Fukano, Mizuki Oshima, Camille Sureau, Noriyuki Watanabe, Takanobu Kato, Asako Murayama, Yingfang Li, Ikuo Shoji, Kunitada Shimotohno, Kazuaki Chayama, Masamichi Muramatsu, Takaji Wakita, Tomoyoshi Nozaki, Hussein H. Aly

**Author notes:** Corresponding Authors: Hussein H. Aly,; Takaji Wakita.

## Abstract

Intracellular transport via microtubule-based dynein and kinesin family motors plays a key role in viral reproduction and transmission. We show here that Kinesin Family Member 4 (KIF4) plays an important role in HBV/HDV infection. We intended to explore host factors impacting the HBV life cycle that can be therapeutically addressed using siRNA library transfection and HBV/NLuc (HBV/NL) reporter virus infection in HepG2-hNTCP C4 cells. KIF4 silencing resulted in a 3-fold reduction in luciferase activity following HBV/NL infection and suppressed both wild-type HBV and HDV infection. Transient KIF4 depletion reduced surface and raised intracellular NTCP (HBV/HDV entry receptor) levels, according to both cellular fractionation and immunofluorescence analysis (IF). Overexpression of wild-type KIF4 but not ATPase-null KIF4 regains the surface localization of NTCP in these cells. Furthermore, immunofluorescence (IF) revealed KIF4 and NTCP colocalization across microtubule filaments, and a co-immunoprecipitation study showed that KIF4 physically binds to NTCP. KIF4 expression is regulated by FOXM1. Interestingly, we discovered that RXR agonists (Bexarotene, and Alitretinoin) down-regulated KIF4 expression via FOXM1-mediated suppression, resulting in a substantial decrease in HBV-Pre-S1 protein attachment to HepG2-hNTCP cell surface and subsequent suppression of HBV infection in HepG2-hNTCP and primary human hepatocytes (PXB) (Bexarotene, IC_50_ 1.89 ± 0.98 μM). Overall, our findings show that human KIF4 is a critical regulator of NTCP surface transport and localization, which is required for NTCP to function as a receptor for HBV/HDV entry. Furthermore, small molecules that suppress or alleviate KIF4 expression would be potential antiviral candidates that target HBV and HDV entry phases.

**Author Summary:** Understanding HBV/HDV entry machinery and the mechanism by which NTCP (HBV/HDV entry receptor) surface expression is regulated is crucial to develop antiviral entry inhibitors. We found that NTCP surface transport is mainly controlled by the motor kinesin KIF4. Surprisingly, KIF4 was negatively regulated by RXR receptors through FOXM1-mediated suppression

This study not only mechanistically correlated the role of RXR receptors in regulating HBV/HDV entry but also suggested a novel approach to develop therapeutic rexinoids for preventing HBV and/or HDV infections in important clinical situations, such as in patients undergoing liver transplantation or those who are at a high risk of HBV infection and unresponsive to HBV vaccination.

## Introduction

Hepatitis B virus (HBV) affects about 250 million individuals globally and is a major cause of chronic liver inflammation. Cirrhosis, liver failure, and liver cancer can all result from a protracted condition of hepatic inflammation and regeneration (1). Sodium taurocholate cotransporting polypeptide (NTCP) was discovered in 2012 to be a key cellular receptor for HBV and its satellite hepatitis delta virus (HDV), which shares the same envelope as HBV (2, 3). When HBV nucleocapsid infects human hepatocytes, it is carried to the nucleus, where the partially double-stranded rcDNA genome is repaired to covalently closed circular (ccc) DNA. This episomal DNA acts as a template for all viral transcripts and pregenomic RNA, forming a very stable minichromosome that is the primary cause of chronic HBV infection, the generation of antiviral escape mutants, or relapse after ceasing nucleo(t)ide analog anti-HBV treatment (4).

Kinesins are a vast protein superfamily that is responsible for the movement of numerous cargos within cells such as membrane organelles, mRNAs, intermediate filaments, and signaling molecules along microtubules (5). Kinesins are also thought to regulate cell division, cell motility, spindle assembly, and chromosomal alignment/segregation (6, 7). KIF4 is a highly conserved member of kinesin family (8–10). KIF4 is also known to move to the nucleus during mitosis, where it interacts with chromatin to alter spindle length and control cytokinesis (11). KIF4A has previously been shown to improve the transport of HIV and adenovirus capsids early in infection (12, 13). As a result, KIF4A might be a promising antiviral target. The transcriptional activator Forkhead box M1 (FOXM1) has been shown to increase KIF4A expression in hepatocellular carcinoma (HCC) (14). Interestingly, HBV upregulates KIF4 expression in HepG2 hepatoma cells, and it was reported to be considerably higher in HBV-associated liver malignancies (15); no information on the role of KIF4 in HBV infection is currently known.

We performed functional siRNA screening using an HBV reporter virus and HepG2-hNTCP cells to uncover host factors that impact the HBV life cycle. We identified KIF4 as a positive regulator for the early phases of HBV/HDV infection based on the findings of this screen. Further investigation indicated that KIF4 is a critical component in the transport and surface localization of NTCP, where it can function as a receptor for HBV/HDV entry. RXR agonists like Bexarotene reduced KIF4 expression and HBV/HDV infection by targeting FOXM1. This is the first study to show that KIF4 plays an essential role in HBV/HDV entry and that it may be used to build effective anti-HBV entry inhibitors.

## Results

### KIF4 is a proviral host factor required for the early stages of HBV infection

We previously used HBV particles containing a chimeric HBV genome (HBV/NL), in which HBV core region is substituted by NanoLuc (NL) gene, to infect HepG2-hNTCP cells formerly transfected with a druggable genome siRNA library two days before HBV/NL infection. We looked at 2,200 human genes to see if they had any effect on the HBV life cycle (16). HBV/NL does not replicate because the HBV core protein (HBc) is not expressed, and the NL levels released after infection only represent the early phases of HBV infection, from the entry through transcription of HBV pregenomic RNA (pgRNA). For each plate, nontargeting or anti-NTCP siRNAs were employed as controls (Fig. 1A). The XTT (2,3-bis-[2-methoxy-4-nitro-5-sulfophenyl]-2H-tetrazolium-5-carboxanilide) test was used to assess cell viability; wells with ≥20% loss of cell viability were removed from further investigation. Previously, we discovered host factors with anti-HBV action (16). In this paper, we describe the discovery of new host factors (proviral factors) necessary in the early stages of HBV infection. The independent silencing of just 14 of the 2,200 host genes (0.6%) reduced NL activity by more than 70% (average of three distinct siRNAs) (Fig. 1B). These genes were identified as pro-HBV host factors (NTCP, SAT1, DVL3, AOX1, DGKH, CAMK1D, CHEK2, AK3, ROCK2, RYK, KIF4, TFIP11, SLC39A6, MXRA5). KIF4 was previously shown to be induced in vitro by HBV expression *in vitro* (15). We also looked at data from an accessible database (17) and discovered that KIF4 was expressed at considerably greater levels in patients with persistent HBV infection than in healthy people (*P* < 0.001). (Fig. 1C). The role of KIF4 in HBV life cycle is not yet reported, hence, we performed further investigation to clarify it. Silencing KIF4 expression with two distinct siRNA sequences (Fig. 1E) led to a 2-fold (*P* < 0.001) or 3-fold (*P* < 0.001) reduction in NL activity relative to cells transfected with the control siRNA (Fig. 1D).

**FIG. 1.**
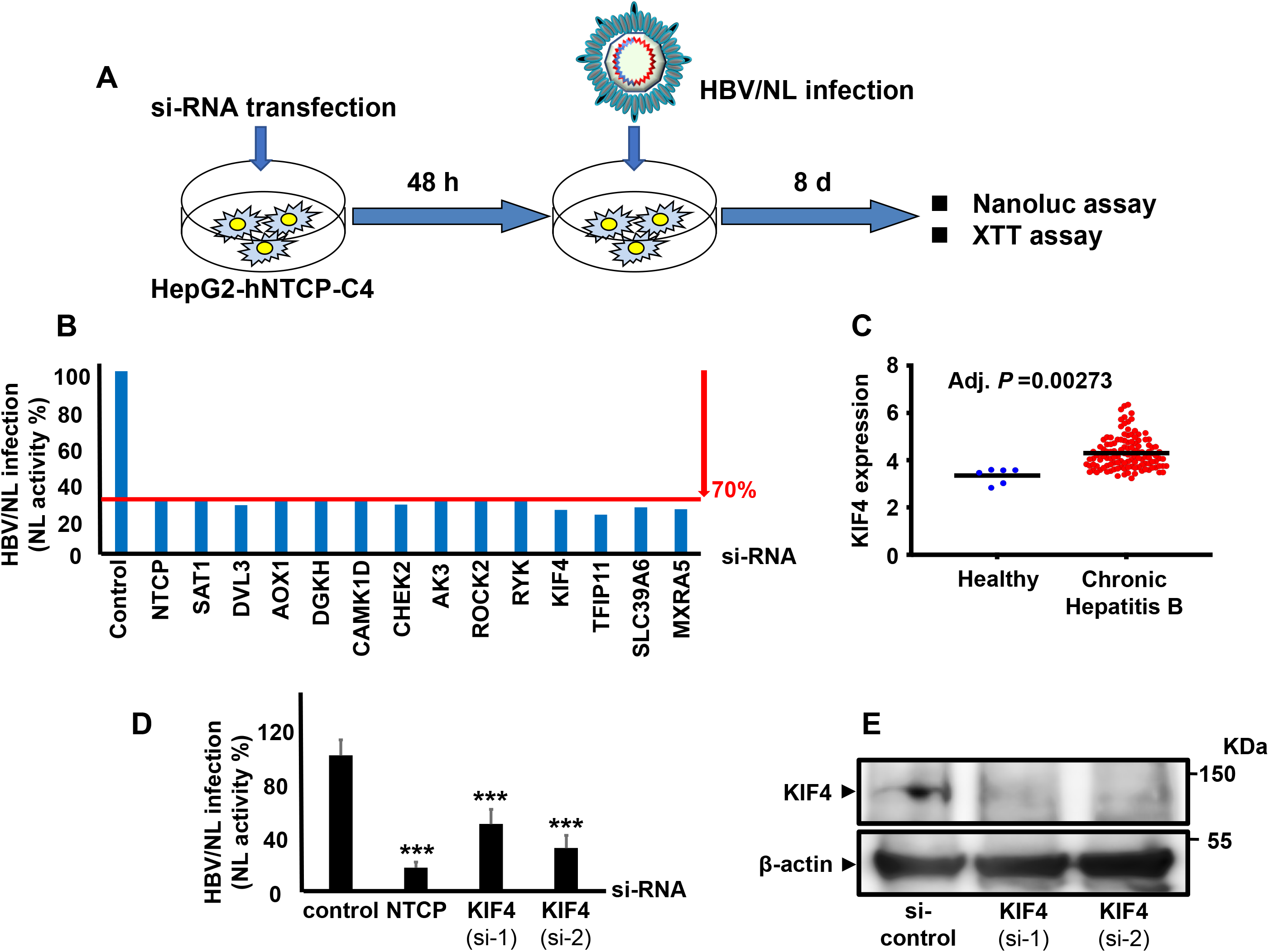
KIF4 is a proviral host factor required for HBV infection and its expression is enhanced in chronic hepatitis B patients. **(A)** A schematic representation of the experimental HBV/NL infection schedule in HepG2-hNTCP used for siRNA library screening. **(B)** HepG2-hNTCP cells were transfected with control, NTCP, or host gene targeting siRNAs from the Silencer Select Human Druggable Genome siRNA Library V4 (Thermo Fisher Scientific, 4397922); host genes siRNA plates were screened as described in the *Material* and *Methods* section. After 2 days of transfection, the cells were inoculated with the HBV/NL reporter virus. At 8 dpi, the luciferase assays were performed, and the NL activity was measured and presented as a percentage relative to control siRNA transfected cells. Of the 2,200 host genes, only 14 genes showed an average of ≥70% reduction of the NL activity upon silencing with a minimum of two independent siRNAs. **(C)** The KIF4 mRNA levels in the liver tissues of patients with chronic HBV infection (n = 122) and healthy subjects (n = 6) (GEO accession number GSE83148). **(D)** HepG2-hNTCP cells were transfected with si-control, si-NTCP, or siRNAs against KIF4 (si-1, and si-2) for 2 days and then inoculated with the HBV/NL reporter virus. At 8 dpi, the cells were lysed, and the luciferase assays were performed, and the NL activity was measured, normalized to cell viability, and plotted as fold changes, relative to control siRNA transfected cells. **(E)** HepG2-hNTCP cells were transfected with control siRNA or siRNAs against KIF4 (si-1 and si-2); the total protein was extracted after 3 days. The expression of endogenous KIF4 (*upper panel*) and β-actin (loading control) (*lower panel*) was analyzed by immunoblotting with the respective antibodies. Statistical significance was determined using Student’s *t*-test (***, *P* < 0.001). For panel (C), statistical significance was evaluated by GEO2R.

### KIF4 is required for cell culture-derived HBV (HBVcc) infection

We used authentic HBV particles generated from grown HepAD38.7-Tet cells to validate the relevance of KIF4 in the HBV life cycle (HBVcc). We reduced KIF4 expression in HepG2-hNTCP cells by transfecting particular siRNA 72 hours before HBVcc infection (Fig. 2A); after 10 to 13 days post infection (pi), we examined its influence on hepatitis B surface antigen (HBsAg), DNA, and HBc levels. As a positive control, si-NTCP was utilized. Silencing KIF4 expression resulted in a substantial decrease of HBsAg in the culture supernatant (*P* < 0.001) to levels equivalent to silencing NTCP expression (Fig. 2B). siKIF4 also decreased overall HBV-DNA levels as measured by southern blot (Fig. 2C) and inhibited HBc expression as measured by immunofluorescence (IF) (Fig. 2D) to levels equivalent to si-NTCP without compromising cellular viability (Fig. 2E). We employed primary human hepatocytes (PXB) cells to examine the effect of suppressing KIF4 expression on HBV infection in a more physiologically relevant paradigm (Fig. 2F). As predicted, inhibiting KIF4 or NTCP expression dramatically reduced HBs levels by ELISA (*P* < 0.01) (Fig. 2G) and decreased extracellular HBV-DNA levels by real-time PCR (*P* < 0.001). (Fig. 2H). Figure 2I depicts the silencing efficiency of KIF4 siRNA in PXB cells.

**FIG 2.**
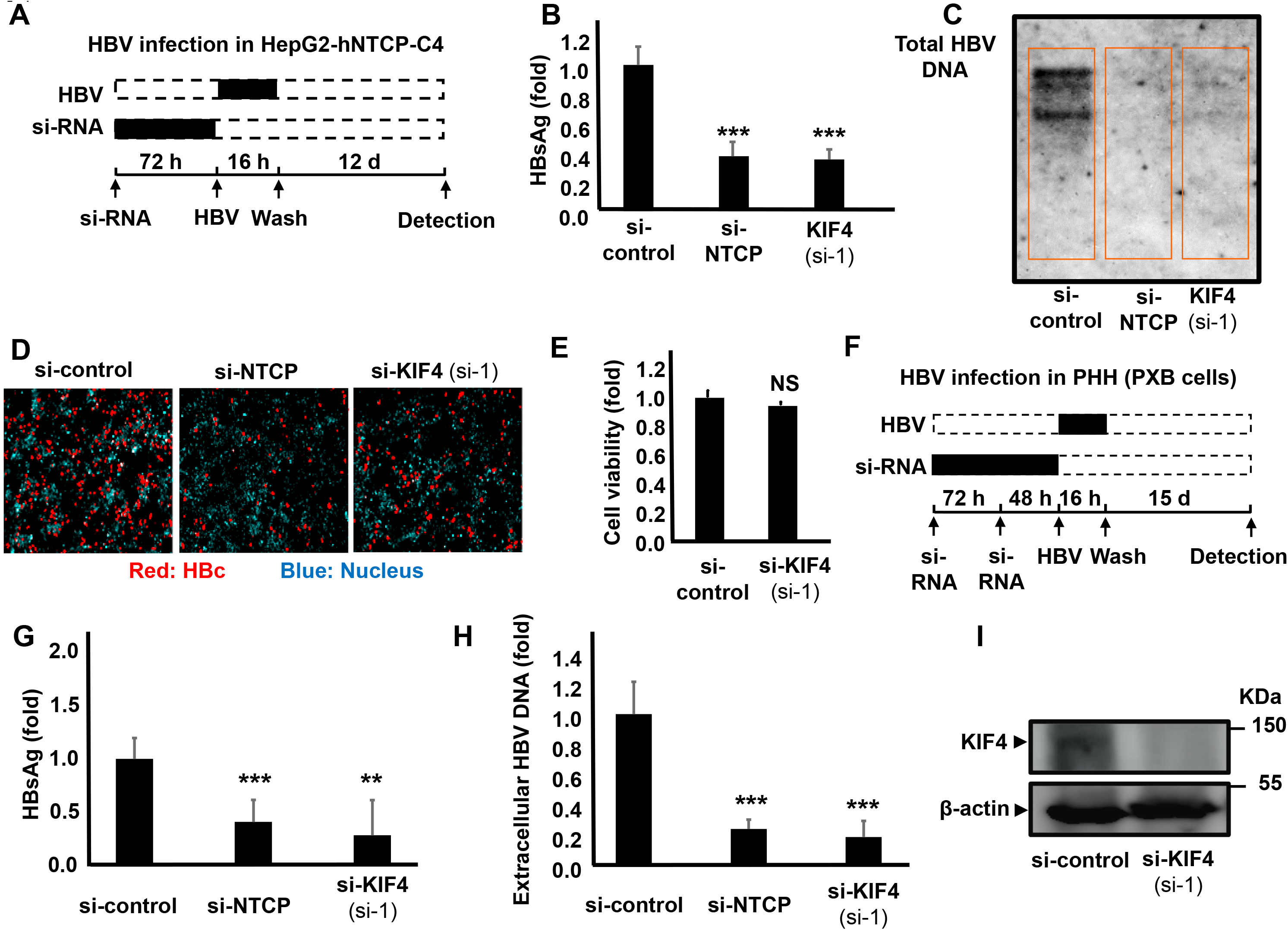
Decreased KIF4 expression suppressed HBV infection in HepG2-hNTCP and primary human hepatocytes (PHH). **(A)** Schematic diagram depicting the scheme for siRNA transfection and subsequent HBV infection in HepG2-hNTCP; HepG2-hNTCP cells were transfected with si-control, si-NTCP, or si-KIF4 (si-1) for 72 h and then inoculated with HBV at 6,000 GEq/cell in the presence of 4% PEG8000 for 16 h. After free HBV were washed out, the cells were cultured for an additional 12 days, followed by the detection of different HBV markers. Black and dashed boxes indicate the interval for treatment and without treatment, respectively. **(B)** HBsAg secreted into the culture supernatant was collected at 10 dpi, quantified by ELISA, and presented as fold changes, relative to the values of control siRNA transfected cells. **(C)** Intracellular HBV DNA and **(D)** HBc protein in the cells were detected at 13 dpi by Southern blot analysis and immunofluorescence, respectively. Red and blue signals in panel **(D)** depict the staining of HBc protein and nucleus, respectively. **(E)** Cell viability was also examined by the XTT assay. **(F)** Schematic diagram showing the scheme for siRNA transfection and the subsequent HBV infection in primary human hepatocytes (PXB); primary human hepatocytes were twice transfected with si-control, si-NTCP, or si-KIF4 (si-1) for consecutive 72 h and 48 h, followed by HBV inoculation at 1,000 GEq/cell in the presence of 4% PEG8000 for 16 h. After being washed, the cells were cultured for an additional 15 days. **(G)** HBsAg and **(H)** Extracellular HBV-DNA secreted into the culture supernatant were quantified by ELISA and real-time PCR, respectively, and the data were presented as fold changes, relative to the values of control siRNA-transfected cells. In all infection assays including siRNA transfection, control siRNA and NTCP-targeting siRNA were used as negative and positive controls, respectively. (**I**) Primary human hepatocytes were twice transfected with siRNAs (as shown in Fig. 2 F); the total protein was extracted and KIF4 (*upper panel*) and β-actin (loading control) (*lower panel*) expression were analyzed by immunoblotting with the respective antibodies. All assays were performed in triplicate and included 3 independent experiments. Standard deviations are also shown as error bars. Statistical significance was determined using Student’s *t-*test (**, *P* < 0.01; ***, *P* < 0.001; NS, not significant).

### KIF4 regulates HBV and HDV entry into host cells

Then, using particular siRNA, we suppressed KIF4 expression and examined the stage of the HBV life cycle that is controlled by KIF4. NTCP plays a role in the specific binding of HBV to the host cell surface by interacting with the preS1 region of HBV’s large surface protein (LHB) (3). We investigated the attachment of a fluorescence-labeled preS1 peptide (6-carboxytetramethylrhodamine-labeled preS1 peptide, or TAMRA-preS1) to HepG2-hNTCP (Fig. 3A). The preS1 binding assay was conducted with si-NTCP as a positive control to validate the specificity of the observed TAMRA-preS1 signals. KIF4 silencing dramatically reduced the interaction between TAMRA-labeled PreS1 and NTCP as identified by IF (Fig. 3A, left pictures) and shown by signal intensities (Fig. 3A, right panel). These findings support the notion that KIF4 is essential for the interaction of HBV and surface NTCP. Using a luciferase reporter system for the various HBV promoters, we discovered that KIF4 expression did not influence the transcriptional activity of these promoters (Fig. 3B). Furthermore, utilizing HBV replicon cells, HepAD38.7-Tet off, which produce HBV pgRNA following tetracycline withdrawal, we discovered that suppressing KIF4 expression did not influence intracellular HBV-DNA levels. (See Fig. 3C.) HDV has the same envelope as HBV and utilizes NTCP as a receptor to enter hepatocytes (2). Consistent with the results obtained in the HBV infection and preS1-binding tests, suppressing KIF4 expression reduced HDV susceptibility in NTCP-expressing cells (Fig. 3D, left pictures), and the magnitude of KIF4 siRNA suppression on HDV infection is displayed in Fig. 3D, right panel. These findings show that KIF4 primarily controlled NTCP-mediated HBV and HDV entry into cells while having little or no influence on HBV transcription or replication.

**FIG. 3.**
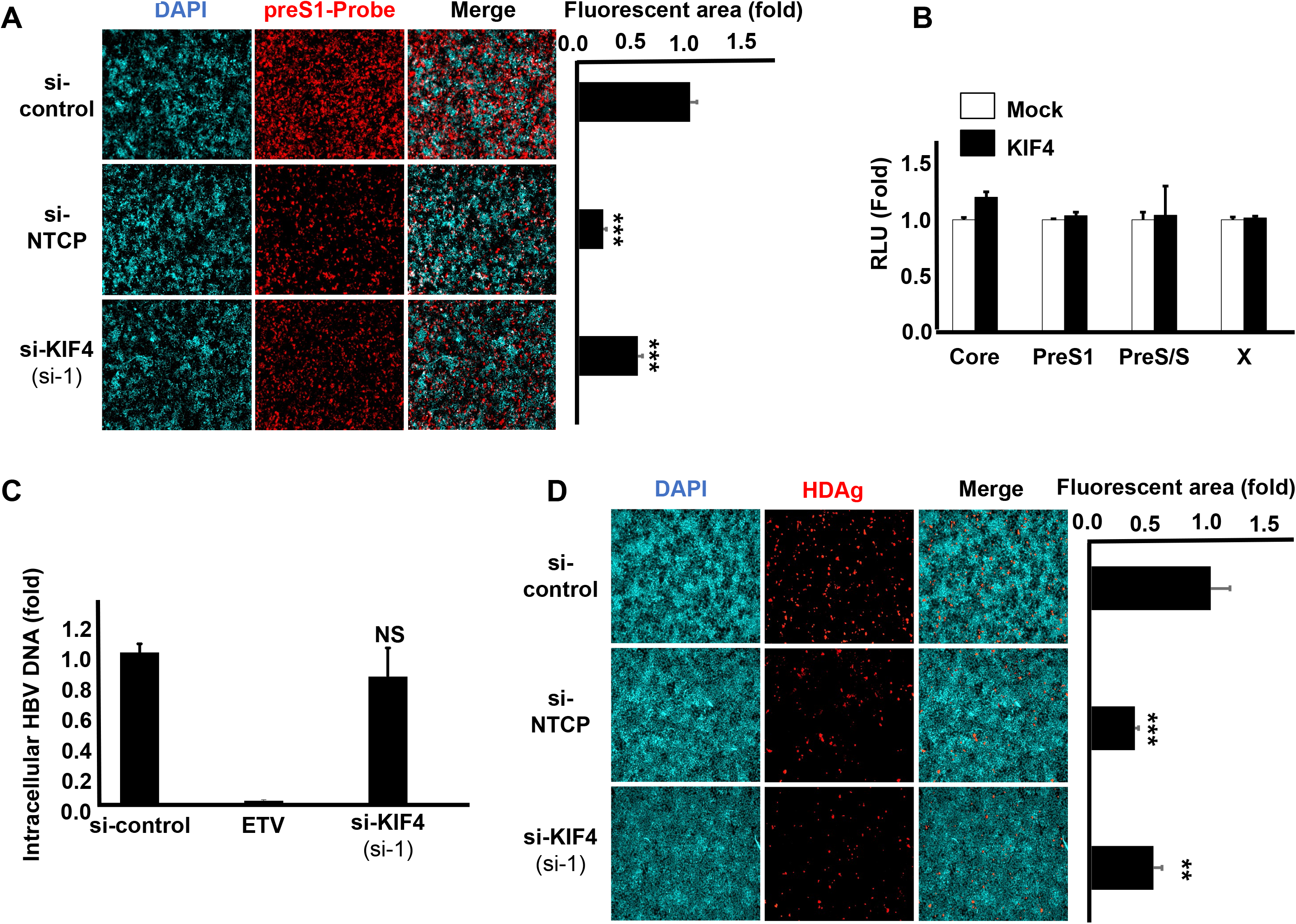
KIF4 knockdown blocks HBV entry and HDV infection into host cells. **(A)** HepG2-hNTCP were transfected with si-control, si-NTCP, or si-KIF4 (si-1) for 72 h and then incubated with 40 nM C-terminally TAMRA labeled and N-terminally myristoylated preS1 peptide (preS1 probe) for 30 min at 37°C (*left panel*); Red and blue signals indicate the preS1 probe and the nucleus, respectively. The fluorescence intensities are shown in the graph (*right panel*). **(B)** HepG2 cells were transfected with a KIF4 expression vector or empty vector (control) together with plasmid vector carrying HBV promoters (Core, X, preS1, or preS2/S) upstream of the *Firefly* luciferase gene and the pRL-TK control plasmid encoding *Renilla* luciferase. At 2 days post-transfection, the cells were lysed and the dual-luciferase activities were measured; the *Firefly* luciferase values were normalized to those of *Renilla* luciferase readings, and the resulting relative luminescence units obtained from KIF4 transfected cells were presented as fold changes compared to the levels detected in the control transfected cells. **(C)** HepAD38.7-Tet cells were transfected with si-control or si-KIF4 (si-1) or treated with 10-µM entecavir as a positive control in the absence of tetracycline to induce HBV replication; At 4 days post-transfection, the cells were lysed and the intracellular HBV DNA was extracted and quantified by real-time PCR. **(D)** HepG2-hNTCP were transfected with siRNAs (as indicated in Fig. 3A), and then inoculated with HDV virions at 50 GEq/cell in the presence of 5% PEG8000 for 16 h; the cells were then washed out to remove the free virus particles and cultured for an additional 6 days, followed by detection of HDAg by IF (*left panel*); Red and blue signals indicate HDAg and nuclear staining, respectively. The fluorescence intensities are shown in the graph (*right panel*). All assays were performed in triplicate and included three independent experiments. The data were pooled to assess the statistical significance. Data are presented as mean ± SD. **, *P* < 0.01; ***, *P* < 0.001; NS, not significant.

### KIF4 regulates surface NTCP expression

We investigated the influence of KIF4 on total and subcellular (both surface and cytoplasmic) NTCP expression after discovering that it is necessary for HBV entrance via regulating the interaction between HBV preS1 and NTCP. Silencing of KIF4 expression did not affect total cellular NTCP protein levels, as shown in figure 4A, but IF examination revealed that silencing of KIF4 disrupted NTCP surface localization and encouraged its accumulation in the cytoplasm (Fig. 4B). This conclusion was supported by biochemical investigation, which indicated that silencing KIF4 dramatically decreased surface NTCP in membranous fraction (Fig. 4C) while increasing intracellular NTCP protein levels in the cytoplasmic fraction (Fig. 4D). Figure 4E depicts band densitometry. It is worth mentioning that silencing KIF4 did not influence surface cadherin or cytoplasmic GAPDH protein levels, indicating that KIF4 has a particular effect on NTCP surface localization.

**FIG 4.**
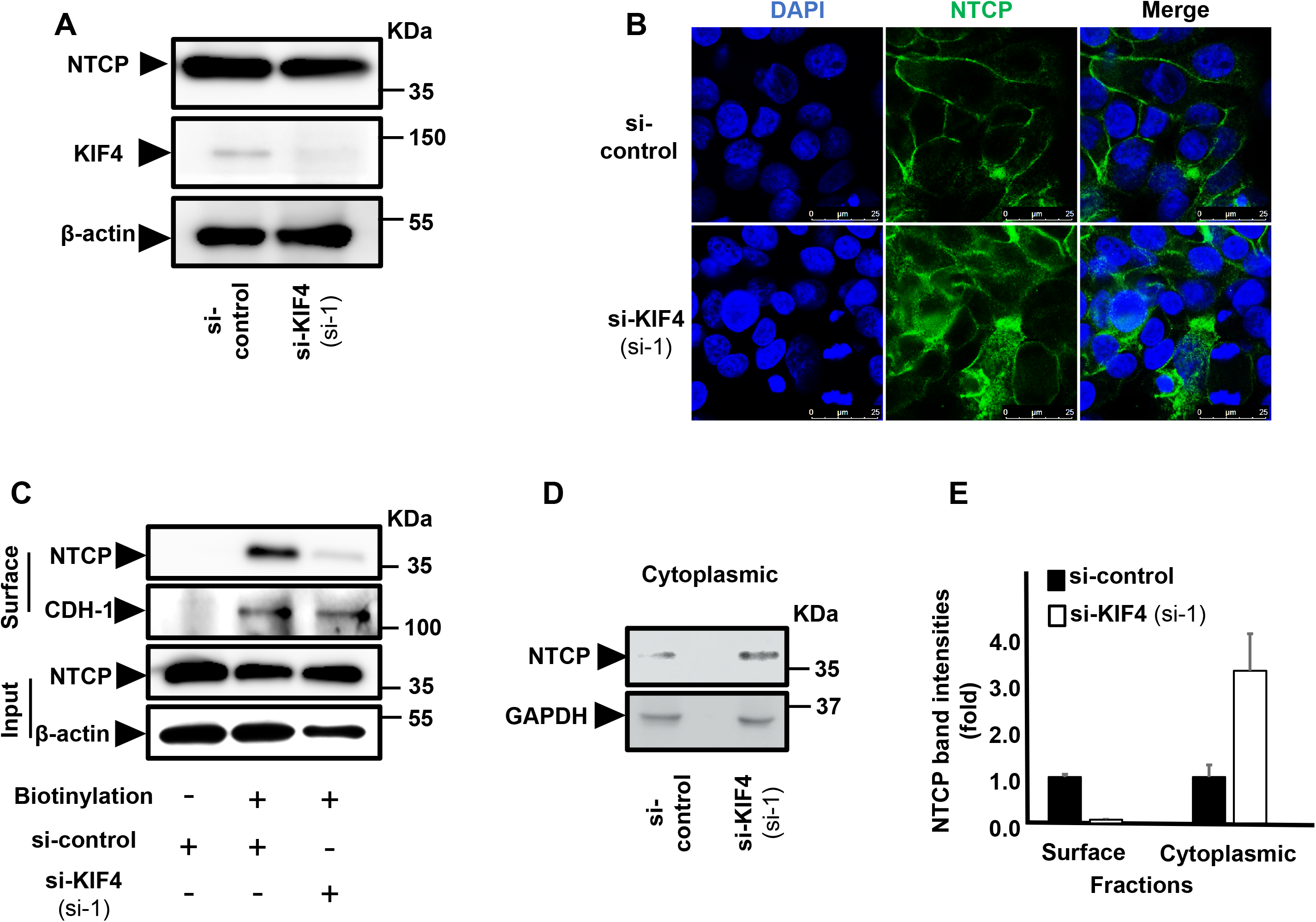
KIF4 regulates the surface NTCP expression. **(A)** HepG2-hNTCP were transfected with either si-control or si-KIF4 (si-1) and incubated for 72 h; the cells were then lysed and the expressions of total NTCP (*upper panel*), KIF4 (*middle panel*), and β-actin (loading control) (*lower panel*) were examined in the whole protein lysate by Western blotting. **(B)** HepG2-hNTCP were transfected with si-RNAs (as in Fig. 4A) and incubated for 72 h; the cells were then fixed with 4% paraformaldehyde, permeabilized with 0.3% Triton X-100, and stained with NTCP antibody and visualized with confocal microscopy. Green and blue signals depict the staining of NTCP (both surface and cytoplasmic), and nuclei, respectively. **(C)** HepG2-hNTCP were transfected with si-RNAs (as in Fig. 4A); at 3 days post-transfection, the cells were surface biotinylated or PBS treated at 4°C for 30 min before cell lysis. After centrifugation and the removal of cell debris, the cell lysates were collected and an aliquot (1/10 volume) was used for the detection of NTCP protein (*Input; upper panel*) and β-actin (loading control) (*Input; lower panels*) by immunoblotting. The remaining cell lysates (9/10 of the original volume) were subjected to pull-down via incubation with pre-washed SA beads for 2 h at 4°C; after washing, the biotinylated surface proteins were eluted and subjected to western blotting in order to detect the surface NTCP and CDH-1 (loading control for surface fraction) (*Surface; upper*, and *lower panels*) with the respective antibodies. **(D)** After transfection with siRNAs (as shown in Fig. 4A), HepG2-hNTCP cells were lysed and the cytoplasmic fraction was isolated, harvested using the Minute ™ Plasma Membrane Protein Isolation and the Cell Fractionation Kit and then subjected to immunoblotting in order to detect the cytoplasmic NTCP (*upper panel*) and GAPDH (loading control for cytoplasmic fraction) (*lower panel*). **(E)** The intensities of both the surface (normalized to CDH-1) and cytoplasmic (normalized to GAPDH) NTCP bands were quantified by ImageJ software and presented as fold changes relative to the control siRNA transfected cells.

### KIF4 motor activity is required for surface NTCP expression

Kinesins, such as KIF4, are motor proteins that hydrolyze ATP to transport different molecules along microtubules (7). Because KIF4 is necessary for surface NTCP expression, we hypothesized that it may function as a transporter that delivers NTCP to the cell surface. As a result, we investigated the function of KIF4 ATPase activity in NTCP surface expression. The sequence of an ATPase-null KIF4 was previously described (18). (Fig. 5A). To mute endogenous KIF4 expression, we utilized a tailored siRNA sequence (si-KIF4 3′ UTR) that targeted the 3′ UTR region of the KIF4 transcript (19) and compensated for this suppression by transfecting plasmids expressing the Myc-tagged WT or ATPase-null KIF4 sequences. Because they lack the 3′ UTR of endogenous KIF4 transcripts, the mRNAs of these constructs are resistant to si-KIF4 3′ UTR. Cellular fractionation revealed that the expression of WT or ATPase-null mutants did not affect surface cadherin (CHD-1) expression, as predicted. Surface NTCP levels are considerably enhanced when KIF4 depletion is compensated with WT, but not ATPase-null KIF4 protein (Fig. 5B). These findings were supported by IF analysis, which revealed significantly greater levels of surface NTCP when endogenous KIF4 silencing was compensated for by WT-KIF4, but not by ATPase-null KIF4 (Fig. 5C, upper panels) (Fig. 5C, lower panels). The si-KIF4 3′ UTR exhibited a substantial reduction of KIF4 by real-time RT-PCR and no cellular cytotoxicity as determined by the XTT assay (Fig. 5D and E). Overall, our findings indicated that KIF4 ATPase (motor) activity is necessary for NTCP surface expression (transport).

**FIG 5.**
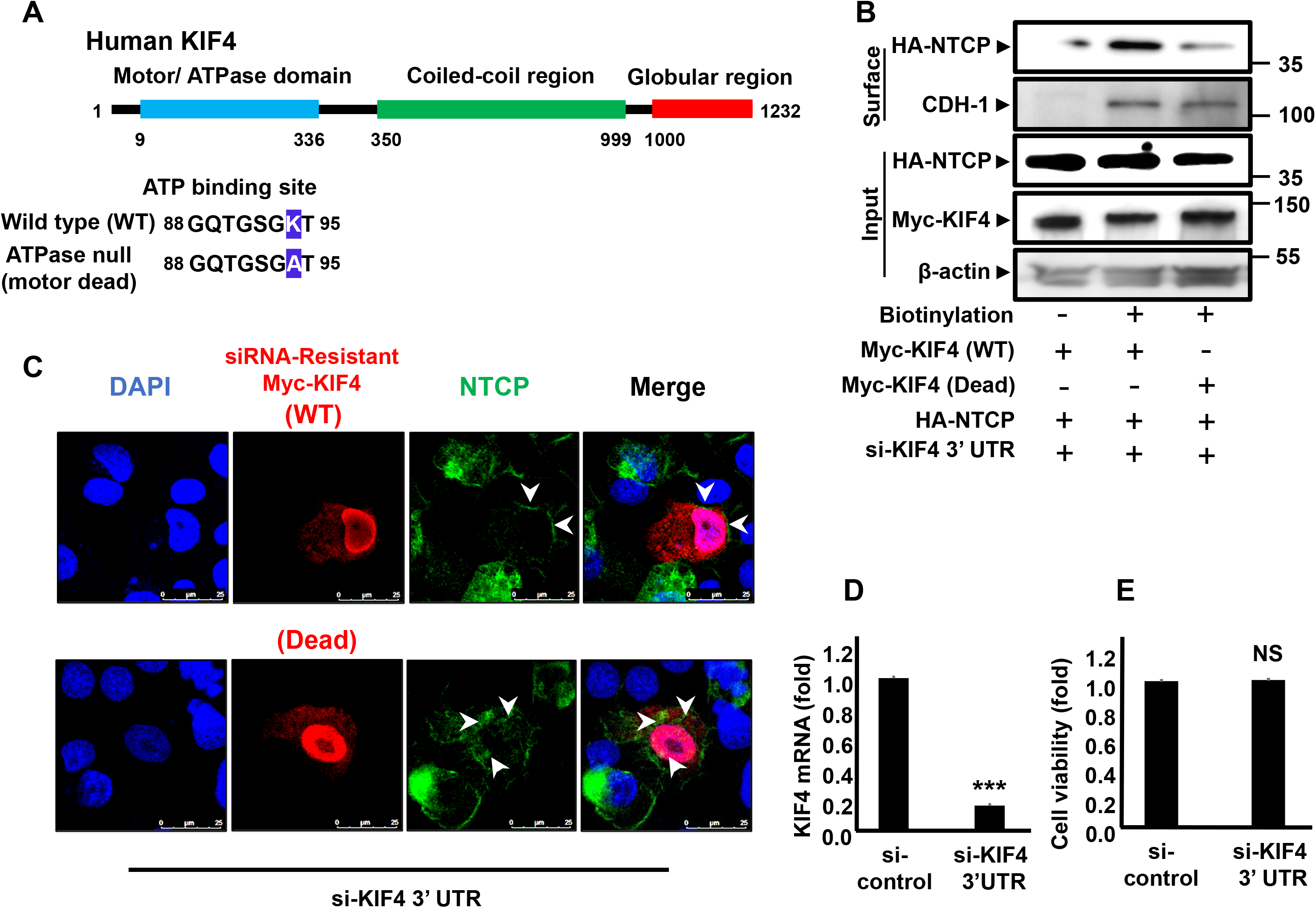
KIF4 motor activity is required for the surface NTCP expression. **(A)** A schematic diagram illustrating human KIF4 domains and the key regions are presented at the top of the figure. Two sequence alignments show ATP-binding Walker A consensus site in the KIF4 motor domain with lysine 94 (Wild type, *upper sequence*) was mutated to alanine (ATPase-null motor dead mutant, *lower sequence*). **(B)** HepG2 cells were transfected with si-KIF4 3′ UTR (targeting endogenous KIF4 mRNA 3’UTR region) along with HA-NTCP, and Myc-KIF4 (WT or ATPase-null) plasmid vectors for 72 h; the cells were then surface biotinylated or PBS treated at 4°C for 30 min before cell lysis. After centrifugation and the removal of cell debris, cell lysates were collected and an aliquot (1/10 volume) was used for the detection of HA-NTCP (*input; upper panel*), Myc-KIF4 (*Input; middle panel*), and β-actin (loading control) (*Input; lower panel*) by immunoblotting. The remaining cell lysates (9/10 of the original volume) were subjected to pull-down via incubation with pre-washed SA beads for 2 h at 4°C; after being washed, the biotinylated surface proteins were eluted and subjected to western blotting in order to detect surface HA-NTCP (*Surface; upper panel*) and CDH-1 (loading control for surface fraction) (*Surface; lower panel*) with the respective antibodies. **(C)** HepG2-hNTCP were transfected with si-KIF4 3′ UTR and Myc-KIF4 WT (*upper panel*) or motor dead mutant (*lower panel*) plasmid vectors; at 3 days post-transfection, the cells were fixed, permeabilized, stained with the indicated antibodies, and visualized by confocal microscopy. Green, red, and blue signals represent the staining of NTCP, Myc-KIF4, and nuclei, respectively. The arrow heads show NTCP localization in the WT or ATPase-null KIF4 transfected cells. **(D)** HepG2-hNTCP were transfected with si-control or si-KIF4 3′ UTR for 48 h; the cells were then lysed and the total RNA content was extracted and the KIF4 expression levels were quantified by RT-qPCR and normalized to the expression of ACTB; or **(E)** the cell viability was examined using XTT assay. Data are presented as fold changes, relative to those of the control siRNA-transfected cells. All assays were performed in triplicate and data from 3 independent experiments were included. The data were pooled to assess the statistical significance. Data are presented as mean ± SD. ***, *P* < 0.001; NS, not significant.

### Physical interaction between KIF4 and NTCP

We hypothesized that direct contact between KIF4 and NTCP across the microtubules is necessary for KIF4 to transport NTCP to the cell surface. We transfected HepG2-hNTCP cells with Halo-tagged KIF4 and examined their intracellular colocalization. Interestingly, IF analysis revealed a significant colocalization between KIF4, NTCP, and α-tubulin (a microtubule marker) (Fig. 6A upper panels). Two distinct cross-sectional lines were constructed (Fig. 6A, center panels), and the colocalization signal intensities were also displayed along these regions of interest (Fig. 6A lower panel). Overlap of KIF4, NTCP, and α-tubulin signal peaks indicated KIF4 and NTCP colocalization along microtubules. The co-immunoprecipitation study verified the direct interaction between KIF4 and NTCP. Myc-tagged KIF4 and HA-tagged NTCP expressing vectors were co-transfected into HEK293-FT cells. We only noticed NTCP co-immunoprecipitation when we used Myc antibody to pull down Myc-tagged KIF4, but not when we used control-IgG (Fig. 6B). Overall, our findings indicate that KIF4 binds to NTCP directly across microtubules in the cytoplasm and uses its ATPase motor domain to transport and transfer NTCP to the cell surface, where it may function as a receptor for HBV and HDV entry.

**FIG 6.**
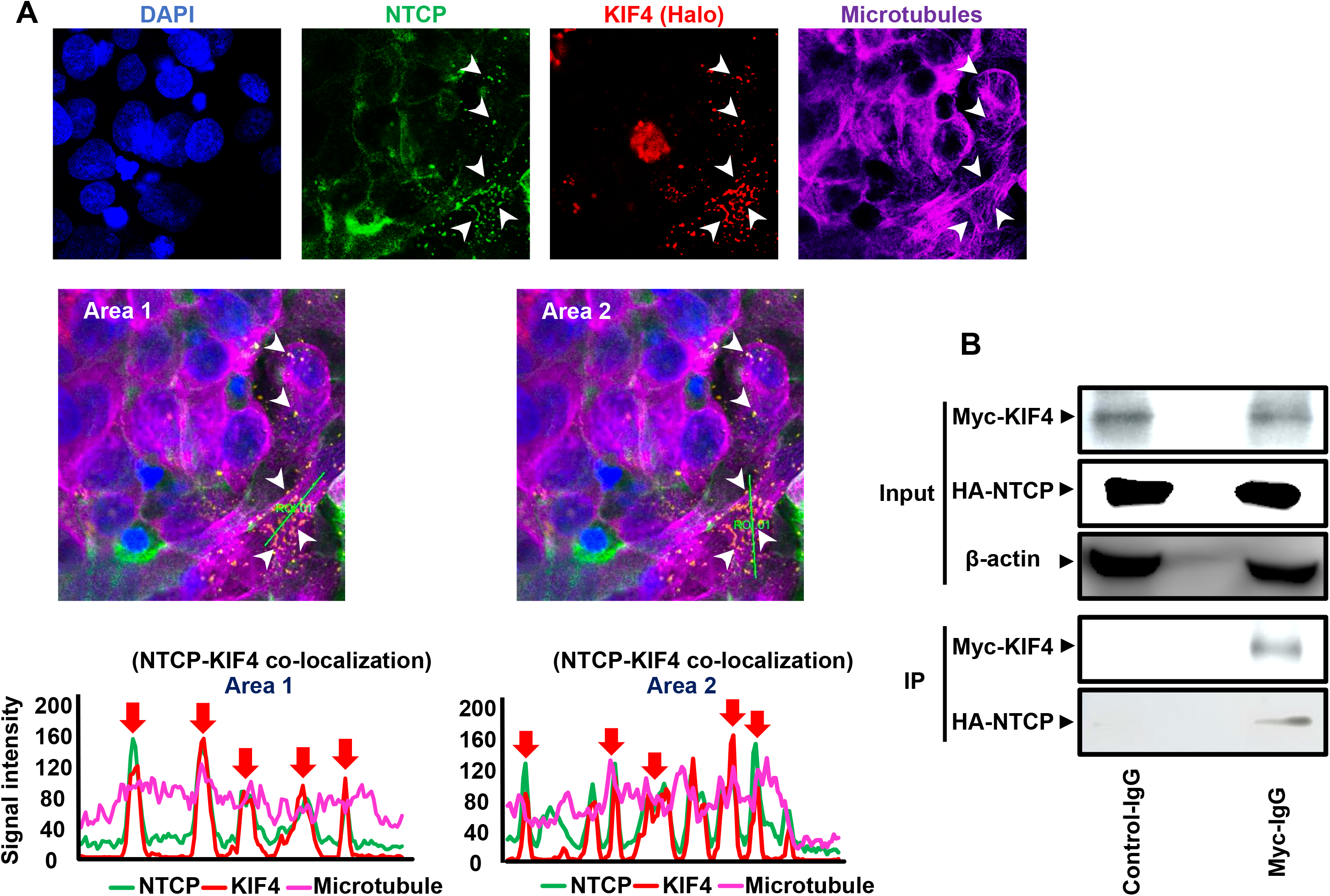
Interaction of KIF4 and NTCP over microtubule filaments. **(A)** HepG2-hNTCP were transfected with a Halo-tagged KIF4 expression vector or empty vector (control). At 48 h post-transfection, the cells were incubated with Halo-tag TMR ligand for 15 min at 37°C; after being washed, cells were fixed, and permeabilized. The cells were stained with antibodies against NTCP and α-tubulin, as indicated in the *Materials* and *Methods* section and examined by confocal microscopy. Blue, green, red, and purple signals indicate nuclear, NTCP, KIF4, and microtubular (α-tubulin) staining, respectively (*upper panel*). White arrows indicate colocalization signals of NTCP and KIF4 over microtubule filaments. The middle panel shows two irrelevant lines crossing different regions of interest within the overlay pattern and represented by their corresponding colocalization signal intensity charts (*lower panel*). **(B)** HEK293-FT cells were co-transfected with HA-NTCP, and Myc-KIF4 plasmid vectors (at a ratio of 1:1). At 3 days post-transfection, the cells were lysed and an aliquot of the cell lysate (1/10 volume) was used for the detection of Myc-KIF4 (*Input; upper panel*), HA-NTCP (*Input; middle panel*), and β-actin (loading control) (*Input; lower panel*) by immunoblotting. The remaining cell lysates (9/10 of the original volume) were subjected to co-IP using either isotype control antibody or anti-Myc IgG to pull down Myc-KIF4. Following IP, each sample was analyzed by immunoblotting for Myc-KIF4 (*IP; upper panel*) and co-immunoprecipitated HA-NTCP (*IP; lower panel*). All assays were performed in triplicate and data from 3 independent experiments were included.

### RXR agonists down-regulate KIF4 expression and block HBV entry by FOXM1 - mediated suppression

The transcription factor forkhead box M1 (FOXM1) has been shown to increase KIF4A expression (14) and is thus anticipated to affect surface NTCP expression and HBV entry into hepatocytes. Furthermore, retinoids including retinoid acid receptor (RAR) and retinoid X receptor (RXR) agonists were reported to decrease FOXM1 expression in human oral squamous cell carcinoma (20) and were anticipated to down-regulate its downstream KIF4 expression and, ultimately, HBV entry. To test this theory, we looked at how various retinoids affected the attachment of TAMRA-labeled preS1 peptide to cell surface NTCP in HepG2-hNTCP cells (Fig. 7A). The preS1 binding assay was done in the presence of NTCP inhibitor, Myrcludex B, as a positive control to confirm the specificity of the observed TAMRA-preS1 signals (21). Interestingly, we discovered that Alitretinoin, a RAR/RXR agonist with potent RXR activity, and Bexarotene, a Pan-RXR agonist, significantly reduced TAMRA-labeled PreS1 binding to NTCP as indicated by IF (Fig. 7A); however, Pan-RAR, ATRA, and RARα-agonist, Tamibarotene, did not affect the preS1 probe binding. These findings indicate that RXR agonists selectively decreased surface NTCP localization and inhibited HBV/NTCP interaction. We then performed a cellular fractionation and found that while treatment with Bexarotene did not affect total NTCP expression (Input, Fig. 7B), it effectively suppressed the level of NTCP protein in the cell surface fraction (Fig. 7B). Bexarotene treatment of HepG2-hNTCP cells resulted in a significant reduction of both FOXM1 and KIF4 expression, supporting our hypothesis (Fig. 7C). Consequently, pretreatment with 10 μM Bexarotene dramatically decreased HBV/NL infection in HepG2-hNTCP cells (*P* < 0.001) (Fig. 7D) without altering cell viability (Fig. 7E).

**FIG 7.**
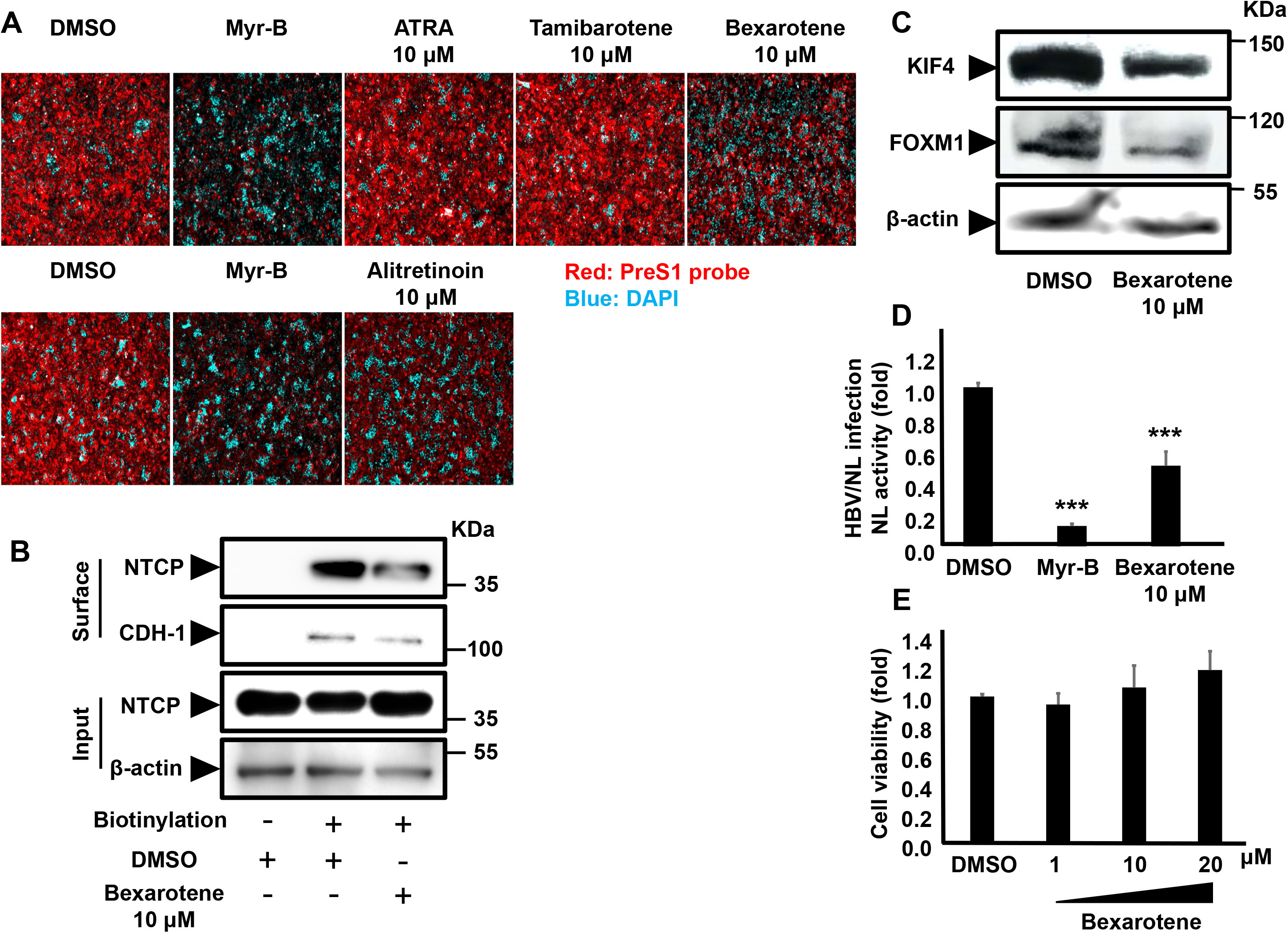
RXR agonists suppressed KIF4 mediated surface NTCP transport, blocked HBV entry, and inhibited HBV/NL infection in HepG2-hNTCP. **(A)** HepG2-hNTCP pretreated with DMSO, 10 μM of the indicated compounds (Bexarotene, ATRA, Tamibarotene) (*upper panel*), or alitretinoin (10 μM) (*lower panel*) for 72 h were incubated with TAMRA-labeled preS1 peptide (preS1 probe) for 30 min at 37°C and then examined by fluorescence microscopy. DMSO-treated cells were incubated with a preS1 probe either in the absence (negative control) or presence (positive control) of 100 nM Myrcludex B (Myr-B); Red and blue signals indicate preS1 probe and the nucleus, respectively. (B) HepG2-hNTCP were treated with DMSO or 10 μM Bexarotene for 72 h, the cells were then surface biotinylated or PBS treated at 4°C for 30 min prior to cell lysis, the cell lysates were collected and an aliquot (1/10 volume) was used for detection of NTCP protein (input) (*Input; upper panel)* and β-actin (loading control) by immunoblotting (*Input; lower panel*). The remaining cell lysates (9/10 of the original volume) were subjected to pull-down via incubation with pre-washed SA beads for 2 h at 4°C; after washing, the biotinylated surface proteins were eluted and subjected to western blotting to detect the surface NTCP (*Surface*; *upper* panel) and CDH-1 (loading control for surface fraction) (*Surface*, *lower panel*) with the respective antibodies. **(C)** HepG2-hNTCP cells treated with DMSO or 10 μM Bexarotene for 72 h were lysed and total cell lysates were subjected to immunoblotting to detect the protein expression levels of KIF4 (*upper panel*), FOXM1 (*middle panel*), and β-actin (loading control) (*lower panel*) with their corresponding antibodies. **(D)** HepG2-hNTCP were pretreated with DMSO or 10 μM Bexarotene for 72 h; then DMSO and Bexarotene were withdrawn from the culture medium 3 h before HBV/NL inoculation, and the cells were inoculated with the HBV/NL reporter virus for 16 h. DMSO-pretreated cells were concomitantly treated with or without 100 nM Myr-B during HBV/NL inoculation. At 8 dpi, the cells were lysed, luciferase assays were performed, and NL activity was measured, and then plotted as fold changes, relative to the values of control DMSO-pretreated cells. **(E)** HepG2-hNTCP were exposed to DMSO or different concentrations of Bexarotene (1 μM, 10 μM, and 20 μM) for 72 h; cell viability was then evaluated by XTT assay. All assays were performed in triplicate and data from 3 independent experiments were included. The data were pooled to assess the statistical significance. Data are presented as mean ± SD. ***, *P* < 0.001.

### Bexarotene pretreatment significantly suppressed HBV and HDV infections in primary human hepatocytes

Bexarotene pretreatment (Fig. 8A) significantly reduced susceptibility to HBV infection in primary human hepatocytes (a more realistic model of HBV infection) in a dose-dependent decrease in secreted HBsAg levels (*P* < 0.001) (Fig. 8B); the 50% inhibitory concentration (IC_50_) was estimated to be 1.89 ± 0.98 μM. Surprisingly, Bexarotene was not harmful to primary human hepatocyte cultures over a wide range of doses, with a 50% cytotoxic concentration (CC_50_) of more than 50 μM. (Fig. 8C). Bexarotene’s selectivity index (CC_50_/IC_50_ ratio) was found to be > 26. We then changed the timing of Bexarotene administration in order to cover the different stages of HBV life cycle in primary human hepatocytes (Bexarotene was administered as follows: pre = 3 days before infection; co = during the inoculation of HBV particles from d0 to d1 pi; post = from d4 to d12 pi; and whole = from 3 days pre infection to d12 pi) (Fig. 8D). The expression of HBc protein by IF was used as a marker of HBV infection. While a modest suppression of HBc detection was found when Bexarotene is added co- or post-infection; the main suppressive effect of Bexarotene on the level of HBc protein was found when it was administered in (pre), or (whole) settings (Fig. 8E [immunofluorescence], and 8F [fluorescence intensity]). There was no apparent difference in Bexarotene-mediated suppression of HBc detection when it was administered in (pre) or (whole) settings. Since the administration of Bexarotene for 3 days before HBV infection is the common time frame between (pre) and (whole) settings, our data suggests that Bexarotene mainly exerts its suppression on HBV when administered before infection. This result is in line with our finding that Bexarotene suppressed surface NTCP localization and subsequent HBV entry; hence its effect is mainly present when administered before (pre) infection. We investigated the effect of Bexarotene pretreatment on HDV infection since HDV has the same surface envelope and so employs NTCP to enter human hepatocytes. Bexarotene reduced HDV infection as predicted, as shown by a decrease in HDV RNA (*P* < 0.001). (Fig. 8G). These findings support the hypothesis that RXR agonists have a suppressive impact on HBV/HDV entry.

**FIG 8.**
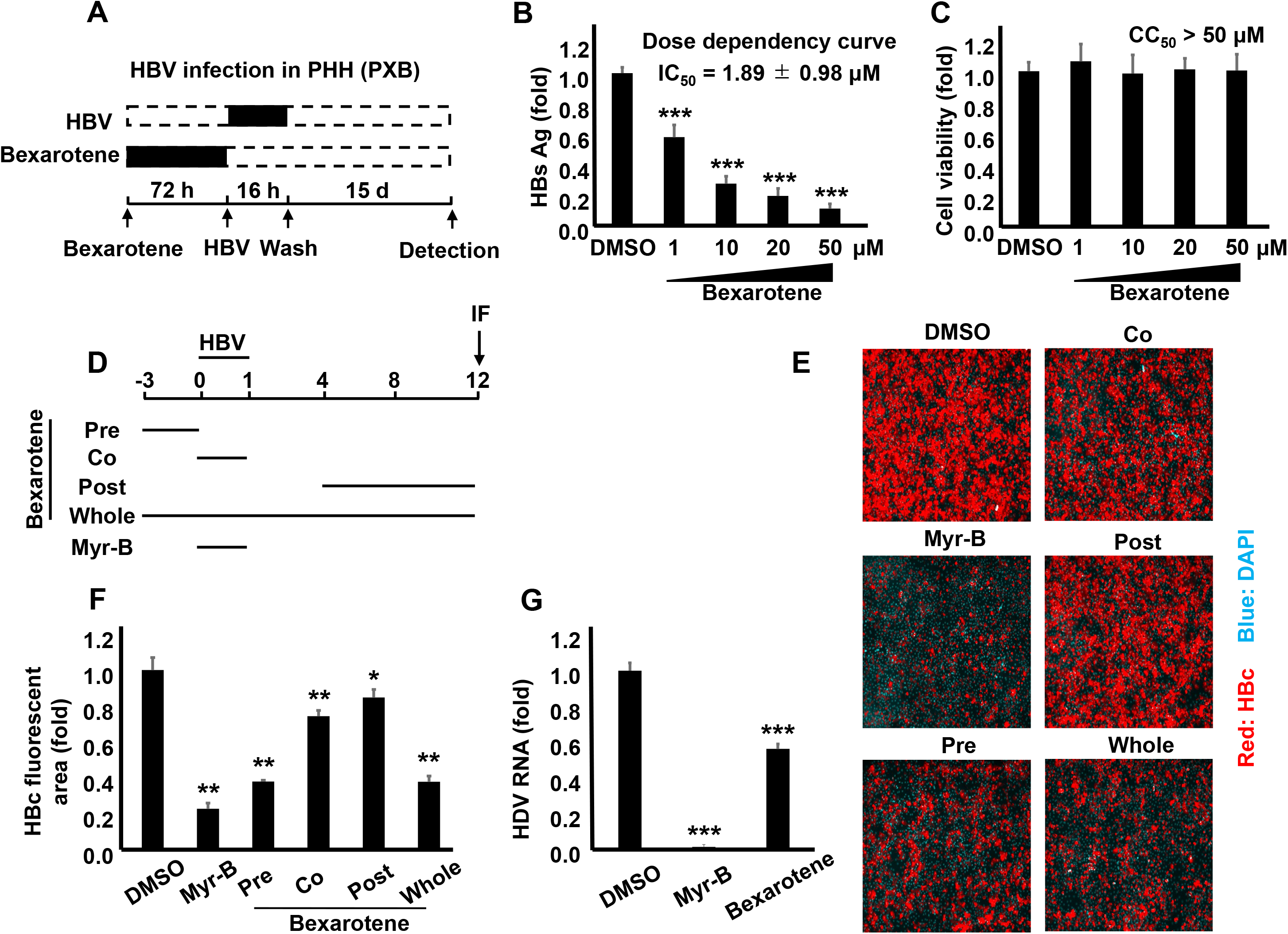
Bexarotene pretreatment significantly suppressed HBV and HDV infections in primary human hepatocytes (PHH). **(A)** A schematic representation showing the protocol used for Bexarotene treatment and subsequent HBV infection in PHH; PHH were pretreated with DMSO or different concentrations of Bexarotene (1 μM, 10 μM, 20 μM, and 50 μM) for 72 h, DMSO and Bexarotene were withdrawn from the culture medium 3 h before HBV infection, and the cells were inoculated with HBV particles at 1,000 GEq/cell in the presence of 4% PEG8000 for 16 h. DMSO-pretreated cells were concomitantly treated with or without 100 nM Myr-B during HBV inoculation. After being washed, the cells were cultured for an additional 15 days. **(B)** HBsAg secreted into the culture supernatant was quantified by ELISA, and the data were presented as fold changes relative to the values of control DMSO-pretreated cells. **(C)** Cell viability were measured by the XTT assay. (**D**) A schematic diagram showing HBV infection protocol; PHH were treated with 15 μM Bexarotene at different time schedule (pre, co, post, and whole) as shown in the figure. The cells were inoculated with HBV at 1,000 GEq per cell in the presence of 4% PEG8000 for 16 h. Bexarotene non-treated cells were concomitantly incubated with or without 100 nM Myr-B during HBV inoculation. After washing out the free virus particles, the cells were cultured for an additional 11 days. (**E**) HBc protein in the cells was detected by immunofluorescence. Red and blue signals depict the staining of HBc protein and nucleus (dapi), respectively and HBc fluorescence intensities are shown in Panel (**F**). **(G)** PHH pretreated with DMSO or Bexarotene (50 μM) for 72 h were inoculated with the HDV at 40 GEq/cell in the presence of 5% PEG8000 for 16 h. DMSO-pretreated cells were concomitantly treated with or without 100 nM Myr-B during HDV inoculation. After washing out the free virus particles, the cells were cultured for an additional 6 days and then lysed; RNA was then extracted and HDV RNA was quantified by RT-qPCR. The data are presented as fold differences relative to those of the control DMSO-pretreated cells. All assays were performed in triplicate and data from 3 independent experiments were included. The data were pooled to assess the statistical significance. For panels **(D and G)**, the assay was performed in triplicate, and data from 2 independent experiments were pooled. Data are presented as mean ± SD. *, *P* < 0.05; **, *P* < 0.01; ***, *P* < 0.001.

## Discussion

The HBV/NL reporter system has previously been described to reflect the early phases of HBV infection (22). We previously reported using this method to screen 2200 druggable human genes and outlined the discovery of MafF and other host factors with anti-HBV function as a result of this screening (16). The current study describes the proviral host factors that are required for the early phases of HBV infection.

KIF4 belongs to the kinesin superfamily (KIFs). KIFs are ATP-dependent microtubule-based motor proteins that are involved in the intracellular transport (11). The N-terminal motor domain of KIF4 is in charge of ATP hydrolysis and microtubule-binding, whereas the C-terminal domain attaches to cargo molecules such as proteins, lipids, and nucleic acids (23). We identified KIF4 as a pro-viral host factor required for the early phases of HBV life cycle. The importance of KIF4 in boosting HBV infectivity early in the HBV life cycle was verified in primary hepatocytes (Fig. 2).

The suppression of the interaction between HBV-PreS1 and HBV entry receptor (surface NTCP) following silencing of KIF4 expression demonstrated the involvement of KIF4 in regulating the function of NTCP as a receptor for HBV entry. HDV and HBV share the same envelope proteins and hence rely on NTCP as an entry receptor into hepatocytes (2). Silencing KIF4 expression dramatically reduced HDV infection, as expected. KIF4 has previously been implicated in the anterograde transport of cellular proteins such as Integrin beta-1 (24), as well as viral proteins such as retroviral (human immunodeficiency virus (HIV-1), murine leukemia virus, Mason-Pfizer monkey virus, and simian immunodeficiency virus) Gag polyprotein (25) to the cell surface to allow for efficient retroviral particle formation. In line with KIF4’s previously described anterograde transport function, we discovered that inhibiting the ATPase motor activity of KIF4 substantially reduced surface and raised cytoplasmic NTCP levels (Fig. 5). We also used immunoprecipitation to validate the physical contact of KIF4 and NTCP, as well as their colocalization on microtubules (Fig. 6). These findings demonstrated that KIF4 controlled the anterograde transport of NTCP to the cell surface, influencing its availability as a receptor for HBV/HDV entry on the hepatocyte surface.

Because the FOXM1 transcription factor is known to influence KIF4 expression (14) it is predicted to affect surface NTCP expression and HBV internalization into hepatocytes. Since retinoids, including RAR and RXR agonists, have been shown to suppress FOXM1 expression in human oral squamous cell carcinoma (20), we hypothesized that retinoid agonists would also suppress KIF4 expression, resulting in impaired transport of NTCP to the cell surface and decreased susceptibility to HBV or HDV infection. Only RXR agonists, Alitretinoin (26), and Bexarotene (27) decreased HBV-PreS1 attachment to NTCP at the cell surface, indicating the specificity of RXR agonists as HBV entry inhibitors (Fig. 7). Bexarotene, a pan-RXR agonist, substantially decreased FOXM1, KIF4, and cell membrane-associated NTCP levels, which in turn inhibited NTCP-dependent HBV-PreS1 binding to the cell surface and the consequent HBV (IC_50_ 1.89 ± 0.98 μM) and/or HDV infection without any evident detrimental impact in primary hepatocytes (Fig. 7) (Fig. 8).

This is the first research to describe Bexarotene as an HBV and HDV NTCP-mediated entry inhibitor. Bexarotene has previously been shown to inhibit the early stages of HBV infection when co-inoculated with HBV during the first 24 hours (28). This impact was influenced in part by RXR-regulated gene expression in arachidonic acid (AA)/eicosanoid biosynthesis pathways, which included the AA synthases phospholipase A2 group IIA (PLA2G2A). The specific step of the HBV life cycle (from attachment to cccDNA formation) impacted by Bexarotene was not defined in that study (28). Furthermore, silencing PLA2G2A expression marginally alleviated Bexarotene’s inhibitory impact on HBV infection (28), indicating the presence of other key Bexarotene-dependent mechanisms that are still inhibiting the early stages of HBV infection. In line with that study, we found that co-treatment of Bexarotene with HBV inoculation moderately suppressed HBV infection, however, we also showed that the major suppression of HBV infection was obtained when Bexarotene was administrated pre-infection (Fig. 8E and F) suggesting the presence of other significant mechanism by which Bexarotene exerts its suppressive effect on HBV infection. Furthermore, we found that Bexarotene administration effectively suppressed surface NTCP levels (Fig. 7B). Hence, our data clearly showed that the main mechanism by which Bexarotene suppressed HBV infection is through the downregulation of surface NTCP levels prior to HBV infection.

Finally, we identified KIF4 as a critical host factor necessary for effective HBV infection. KIF4 controls the levels of surface NTCP by anterograde transport of NTCP to the cell surface, which is needed for NTCP to function as a receptor for HBV and/or HDV entry. NTCP is the major transporter of conjugated bile salts from the plasma compartment into the hepatocyte. Although the loss of surface NTCP expression in a patient with the homozygous SLC10A1 gene containing a R252H point mutation showed higher levels of bile salts in the plasma, however, it did not show any evidence of cholestatic jaundice, pruritis, or liver dysfunction. Importantly, the presence of secondary bile salts in his circulation suggested residual enterohepatic cycling of bile salts (29). Furthermore, in NTCP knockout mice, some showed a reduced body weight, however, most animals showed no signs of cholestasis, inflammation, or hepatocellular damage (30). While further in-vivo data are still required to assess efficiency and safety of NTCP targeting drugs, the available data suggest its possible use for the prophylactic treatment against HBV infection. HBIG is used to inhibit HBV vertical transmission (31). Furthermore, extended therapy with HBIG in conjunction with a nucleos(t)ide analog is necessary following liver transplantation (LT) to reduce the HBV recurrence rate to less than 10% in 1-2 years post-transplantation (32). Bexarotene, which inhibits HBV cell entry, might be utilized as an alternative to HBIG. Because HBV entry inhibition reduces the intrahepatic cccDNA pool (33), entry inhibitors are expected to be useful in preventing de novo infection in clinical settings such as vertical transmission and HBV recurrence post-LT. These data strongly suggest that Bexarotene and its derivatives would be studied further for the development of a new class of anti-HBV agents.

## Materials and Methods

### Cell culture

HepG2, HepG2-hNTCP-C4, HepAD38.7-Tet, primary human hepatocytes (Phoenixbio; PXB cells), and HEK 293FT cells were cultured as previously described (16). For maintenance, HepG2-hNTCP cells were cultured in 400 µg/mL G418 (34), while HepAD38.7-Tet cells were cultured in 0.4 μg/mL tetracycline that is withdrawn from the medium upon induction of HBV replication (35).

### Reagents and compounds

Sulfo-NHS-LC-Biotin (A39257) was purchased from Invitrogen. Myrcludex-B was provided by Dr. Stephan Urban, at the University Hospital Heidelberg. Bexarotene (SML0282), ATRA (R2625), Tamibarotene (T3205), Alitretinoin (R4643), Entecavir, and DMSO were all purchased from Sigma-Aldrich.

### Human genome siRNA library screening

siRNA screening was performed as reported previously (36). Briefly, HBV host factors were screened using the Silencer Select™ Human Druggable Genome siRNA Library V4 transfection in HepG2-hNTCP cells. siRNAs were arrayed in 96-well-plates, and negative control siRNA and si-NTCP were added to control the data obtained from each of the 96-well-plates. siRNAs with different sequences targeting the same genes were distributed across 3 plates (A, B, and C). Plates utilized in this screening are described elsewhere (16).

### HBV/NL preparation and infection assay

Reporter HBV/NL particles carrying recombinant HBV virus encoding NL gene were collected from the supernatant of HepG2 cells transfected by pUC1.2xHBV/NL plasmid expressing HBV genome (genotype C) in which the core region is substituted with NL-encoding gene, and pUC1.2xHBV-D helper plasmid carrying packaging-deficient HBV genome as described previously (22, 36). HBV/NL infection was performed 2 days after siRNA transfection. At 8 dpi, the cells were lysed, and the Nano-Luc reading was measured using the Nano-Glo® Luciferase Assay System (Promega, N1150), according to the manufacturer’s instructions.

### RNA and DNA transfection

The cells were reverse transfected with siRNAs using Lipofectamine RNAiMAX (Invitrogen) according to the manufacturer’s guidelines. Forward siRNA transfection in PXB cells was also performed using Lipofectamine RNAiMAX. Transfection with plasmid DNA was performed using the Lipofectamine 3000 (for HepG2 and HepG2-hNTCP) or Lipofectamine 2000 (For 293FT cells), according to the manufacturer’s protocol. siRNA/Plasmid DNA co-transfection in HepG2 or HepG2-hNTCP was implemented with Lipofectamine 2000.

### Plasmids and siRNAs

N-terminal HaloTag KIF4 expressing plasmid (pFN21ASDB3041) was purchased from Promega. The N-terminal Myc-tagged KIF4 (both wild type and ATPase-null motor inactive mutant) cloned in the pIRESpuro3 expression vector was kindly provided by Dr. Toru Hirota at Cancer Institute of the Japanese Foundation for Cancer Research (JFCR). Myc-tagged KIF4 motor inactive mutant was created by substituting 94 aa lysine in the ATP-binding Walker A consensus site to alanine (37). HA-tagged NTCP was kindly provided by Dr. Hiroyuki Miyoshi at RIKEN BioResource Research Center, Japan (38). pUC1.2xHBV/NL and pUC1.2xHBV-D plasmids were kindly supplied by Dr. Kunitada Shimotohno at National Center for Global Health and Medicine, Japan. pSVLD3 plasmid was kindly provided by Dr. John Taylor at the Fox Chase Cancer Center, USA. Silencer Select™ si-KIF4 (si-1, s24406; si-2, s24408), si-NTCP (s224646), control siRNA (#1), and customized si-KIF4 3′ UTR targeting endogenous KIF4 mRNA 3’-UTR region (5′-GGAAUGAGGUUGUGAUCUUTT-3′) were purchased from Thermo Fisher Scientific.

### HBV infection assay

HBV (genotype D) particles were concentrated from the culture supernatant of HepAD38.7 Tet cells as described elsewhere (34). HepG2-hNTCP and primary human hepatocytes (PXB) were inoculated with HBV at 6000 and 1000 genome equivalent (GEq)/cell, respectively, as described previously (16).

### HBV preS1 binding assay

HBV preS1 peptide spanning 2–48 amino acids of the preS1 region with N-terminal myristoylation, and C-terminal 6-carboxytetramethylrhodamine (TAMRA) conjugation (preS1 probe) was synthesized by Scrum, Inc. EZ-Link™. The attachment of HBV preS1 peptide to the HepG2-hNTCP cell surface was performed and analyzed as described previously (38).

### HDV infection assay

HDV used in the infection assay was derived from the culture supernatant of Huh7 cells co-transfected with pSVLD3 and pT7HB2.7 as previously reported (39, 40). HepG2-hNTCP and primary human hepatocytes (PXB) were infected with HDV at 40–50 GEq/cell as described previously (41).

### Dual-luciferase reporter assay

HepG2 cells were co-transfected with effector plasmid (Mock or KIF4), the *Firefly* luciferase reporter vectors, and the *Renilla* luciferase plasmid pRL-TK (Promega) as an internal control. The reporter plasmids carrying the entire core promoter (nucleotide [nt] 900–1817), preS1 promoter (nt 2707–2847), preS2/S promoter (nt 2937–3204), or Enh1/X promoter (nt 950–1373) upstream of the *Firefly* luciferase gene, has been reported previously (42). At 2 days after transfection, the cells were lysed and the luciferase activities were measured using the GloMax® 96 Microplate Luminometer (Promega, GMJ96).

### HBV replication assay

In the absence of tetracycline, HepAD38.7-Tet cells were reverse transfected with si-control or si-KIF4 or treated with 10 µM entecavir as a positive control. At 4 days post-transfection, the cells were lysed and the intracellular HBV DNA was extracted and quantified by real-time PCR (35).

### Indirect immunofluorescence assay

Immunofluorescence assay was basically performed as described previously (43). primary antibodies used in the study included rabbit anti-HBc (Neomarkers, RB-1413-A), anti-HDAg, anti-NTCP (Sigma, HPA042727), mouse anti-c-Myc (Santa Cruz, sc-40), and anti-α-tubulin (Sigma, T5168). Alexa Flour555-, Alexa Flour488-, or Alexa Flour647-conjugated secondary antibodies (Invitrogen) were utilized together with DAPI to visualize the nucleus. For Halo tag, live cells were treated with cell-permeant Halotag TMR ligand (Promega, G8251) before paraformaldehyde fixation. Microscopic examination of the infected cells or preS1 binding was performed by fluorescence microscopy (KEYENCE, BZ-X710); the observation of the subcellular localization was performed using a high-resolution confocal microscope (Leica, TCS 159 SP8) as described previously (43).

### Immunoblot assay

Immunoblotting and protein detection were essentially performed as previously described (44). Protein detection was performed using the following primary antibodies; mouse monoclonal E-cadherin antibody (Santa-Cruz, sc-8426), anti-GAPDH (Abcam, ab9484), anti-Myc (Santa Cruz, sc-40), anti-β-actin (Sigma-Aldrich, A5441), anti-FOXM1 (Santa Cruz, sc-271746); and rabbit polyclonal anti-KIF4A (Invitrogen, PA5-30492), anti-NTCP (Sigma, HPA042727), anti-HA (Sigma, H6908). For immunoblotting of free or tagged NTCP, the sample was treated with 250-U Peptide-N-Glycosidase F (PNGase F) to digest N-linked oligosaccharides from glycoproteins before loading to SDS-PAGE (43).

### Cell surface biotinylation and extraction of surface proteins

Cell surface biotinylation was performed to separate the surface proteins with streptavidin beads. The cells were washed with PBS and then incubated with 0.5 mg/mL EZ-Link™ Sulfo-NHS-LC-Biotin for 30 min at 4°C to biotinylate the cell surface proteins. After quenching with PBS containing 0.1% BSA and washing with PBS thrice to remove free inactive biotin, the cells were lysed in lysis buffer (150 mM NaCl, 50 mM Tris-HCL PH 7.4, 5 mM EDTA, 1% NP40) containing 1x protease inhibitor (Roche) for 15 min at 4°C. The cell lysate was centrifuged, and the supernatant was harvested and added to pre-washed streptavidin agarose (SA) beads and incubated for 2 h at 4°C (Pull-down step). Finally, the SA beads were washed with lysis buffer, and the adsorbed proteins were eluted in the sample buffer and subjected to immunoblot assay as described earlier. The surface adhesion protein E-cadherin (CDH-1) was used as a loading control for biotinylated surface fraction, as reported elsewhere (45).

### Purification of cytoplasmic fraction

The cells were washed with cold PBS, lysed, and subjected to cell fractionation; the cytosolic fraction was isolated from the whole cell lysate using the Minute ™ Plasma Membrane Protein Isolation and Cell Fractionation Kit (Invent Biotechnologies) according to the manufacturer’s protocol (46).

### Co-Immunoprecipitation (Co-IP) assay

293FT cells were transfected with HA-tagged NTCP and Myc-tagged KIF4 expression plasmids at a 1:1 ratio for the assessment of the possible physical interaction between NTCP and KIF4. At 72 h after transfection, the cells were lysed and subjected to immunoprecipitation with the mouse monoclonal anti-Myc (Santa-Cruz) antibody or mouse normal IgG as a negative control. Cell lysis and Co-IP were conducted using the Pierce™ Co-IP Kit (Thermo Fisher Scientific, 26149) according to the manufacturer’s instructions.

### DNA and RNA extraction

Intracellular HBV DNA was extracted from the cells using the QIAamp Mini Kit (QIAGEN), and the extracellular HBV DNA was recovered from the supernatant using the SideStep Lysis and Stabilization Buffer (Agilent Technologies, 400900), while RNA extraction was performed using the NucleoSpin® RNA XS Kit (MACHEREY-NAGEL) according to the manufacturer’s protocols.

### Southern blot analysis

Southern blotting was performed to detect intracellular HBV DNAs as described previously (43).

### qPCR and RT-qPCR

Real-time PCR (for the detection of total HBV DNA) and reverse transcription real-time PCR (for the measurement of HDV RNA) were essentially performed as previously described (41, 47) using the primer-probe sets; 5′-AAGGTAGGAGCTGGAGCATTCG-3′, 5′-AGGCGGATTTGCTGGCAAAG-3′, 5′-FAM-AGCCCTCAGGCTCAGGGCATAC-TAMRA-3′ for HBV DNA, and 5′-GGACCCCTTCAGCGAACA-3′, 5′-CCTAGCATCTCCTCCTATCGCTAT-3′, 5′-FAM-AGGCGCTTCGAGCGGTAGGAGTAAGA-TAMRA-3′ for HDV RNA. qPCR for Intracellular HBV DNA was performed by the 2^(−ΔΔCT)^ method using chromosomal GAPDH DNA sequence (via primer-probe set Hs04420697_g1; Applied Biosystems) as an internal normalization control. Isolated RNA was reverse-transcribed using the High-Capacity cDNA Reverse Transcription Kit (Thermo Fisher Scientific), and the relative levels of the KIF4 mRNA were determined using the TaqMan Gene Expression Assay with the primer-probe set Hs00602211_g1 (Applied Biosystems), while the ACTB expression (primer-probe set 748 Hs99999903_m1) was included as an internal control for normalization (36).

### ELISA

Cell supernatants were harvested and ELISA quantification of the secreted HBs was performed as described previously (47). The half-maximal inhibitory concentration (IC_50_) value for Bexarotene was calculated as described previously (38).

### Cell viability assay

Cell viability was evaluated using the Cell Proliferation Kit II (XTT) according to the manufacturer’s guidelines (43).

### Database

Transcriptional profiling of patients with chronic HBV (NCBI Gene Expression Omnibus [GEO] accession number GSE83148) was identified in the GEO public database. The expression data for KIF4 were extracted by GEO2R.

### Statistical analysis

Unless mentioned otherwise, the experiments were performed in triplicates, and the means of data from three independent experiments were calculated and presented in mean ± SD. Statistical significance was determined using Two-tailed unpaired student’s *t-*tests (*, *P* < 0.05; **, *P* < 0.01; ***, *P* < 0.001; NS, not significant). For the KIF4 expression level in chronic HBV (NCBI [GEO] accession number GSE83148), statistical significance was evaluated by GEO2R to calculate the adjusted *P* value.

## Acknowledgments

S.A.G was the recipient of the Egyptian Japanese Education Partnership-3 (EJEP-3) PhD scholarship provided by the Ministry of Higher Education of Egypt. This study was supported by a Grant-In-Aid for Scientific Research (19K07586), and grants from the Research Program on Hepatitis from the Japan Agency for Medical Research and Development (AMED; 21fk0310104j0905, 21fk0310109j0405, 21fk0310103j0305, and 20fk0310109h0004). We gratefully acknowledge Dr. Stephan Urban at University Hospital Heidelberg for providing Myrcludex-B; Dr. Toru Hirota at JFCR, Japan for providing Myc-tagged KIF4 (both Wild Type and motor inactive mutant); Dr. John Taylor at the Fox Chase Cancer Center, the USA for providing pSVLD3 plasmid; and Dr. Hiroyuki Miyoshi at RIKEN, Japan, for providing HA-tagged NTCP.

